# Data-based, synthesis-driven: setting the agenda for computational ecology

**DOI:** 10.1101/150128

**Authors:** Timothée Poisot, Richard Labrie, Erin Larson, Anastasia Rahlin

**Author notes:** Correspondence to Timothée Poisot –.

## Abstract

Computational thinking is the integration of algorithms, software, and data, to solve general questions in a field. Computation ecology has the potential to transform the way ecologists think about the integration of data and models. As the practice is gaining prominence as a way to conduct ecological research, it is important to reflect on what its agenda could be, and how it fits within the broader landscape of ecological research. In this contribution, we suggest areas in which empirical ecologists, modellers, and the emerging community of computational ecologists could engage in a constructive dialogue to build on one another’s expertise; specifically, about the need to make predictions from models actionable, about the best standards to represent ecological data, and about the proper ways to credit data collection and data reuse. We discuss how training can be amended to improve computational literacy.

---

Computational science happens when algorithms, software, data management practices, and advanced research computing are put in interaction with the explicit goal of solving “complex” problems. Typically, problems are considered complex when they cannot be solved appropriately with mathematical modelling (i.e. the application of mathematical models that are not explicitly grounded into empirical data) or data-collection only (Dorner & Funke 2017). Computational science is the application to research questions of computational thinking (Papert 1996), i.e. the feedback loop of abstracting a problem to its core mechanisms, expressing a solution in a way that can be automated, and using interactions between simulations and data to refine the original problem or suggest new knowledge. Computational approaches are commonplace in most areas of biology, to the point where one would almost be confident that they represent a viable career path (Bourne 2011). Collecting ecological data is a time-consuming, costly, and demanding project; in addition, the variability of these data is high (both in terms of variance and in terms of quantity and completeness). In parallel, many ecological problems lack appropriate formal mathematical formulations, which we need in order to construct strong, testable hypotheses. For these reasons, computational approaches hold great possibilities, notably to further ecological synthesis and assist decision-making (Petrovskii & Petrovskaya 2012).

Levin (2012) suggested that ecology (and evolutionary biology) should continue their move towards a *marriage of theory and data.* In addition to the lack of adequately expressed models, this effort is hampered by the fact that data and models are often developed by different groups of scientists, and reconciling both can be difficult. This has been suggested as one of the reasons for why theoretical papers (defined as *papers with at least one equation in the main text)* experience a lower number of citations (Fawcett & Higginson 2012); this is the tragic sign that empirical scientists either do not see the value of theoretical work, or have not received the training to usefully rely on math-heavy theoretical papers, which of course can be blamed on both parties. One of the leading textbooks on mathematical models in ecology and evolution (Otto & Day 2007) is more focused with algebra and calculus, and not with the integration of models with data. Other manuals that cover the integration of models and data tend to lean more towards statistical models (Bolker 2008; Soetaert & Herman 2008). This paints a picture of ecology as a field in which dynamical models and empirical data do not interact much, and instead the literature develops in silos.

Computational ecology is the application of computational thinking to ecological problems. This defines three core characteristics of computational ecology. First, it recognizes ecological systems as complex and adaptive; this places a great emphasis on mathematical tools that can handle, or even require, a certain degree of stochasticity to accommodate or emulate what is found in nature (Zhang 2010, 2012). Second, it understands that data are the final arbiter of any simulation or model (Petrovskii & Petrovskaya 2012); this favours the use of data-driven approaches and analyses (Beaumont 2010). On this point, computational approaches differ greatly from the production of theoretical models able to stands on their own *(i.e.* with no data input). Finally, it accepts that some ecological systems are too complex to be formulated in mathematical or programmatic terms (Pascual 2005); the use of conceptual, or “toy” models, as long as they can be confronted to empirical data, is preferable to “abusing” mathematics by describing the wrong mechanism well (May 2004). By contrast, modelling approaches are by construction limited to problems that can be expressed in mathematical terms. To summarize, we define computational ecology as the sub-field tasked with integrating real-world data with mathematical, conceptual, and numerical models (if possible by deeply coupling them), in order to assist with the most needed goal of improving the predictive accuracy of ecological research (Houlahan et al. 2017; Maris et al. 2017). Jorgensen (2008) identified that one of the current challenges is to facilitate the integration of existing data in the explosion of modelling techniques (most of which were designed to answer long-standing questions in ecological research).

Ecology as a whole (and community ecology in particular) circumvented the problem of model and data mismatch by investing in the development and refinement of statistical models (see Warton et al. 2014 for an excellent overview) and “numerical” approaches (Legendre & Legendre 1998) based on multivariate statistics. These models are able to *explain* data, but very rarely do they give rise to new predictions despite it being a very clear priority even if we “simply” seek to further our understanding (Houlahan et al. 2017). Computational ecology can fill this niche; at the cost of a higher degree of abstraction, its integration of data and generative models (*i.e.* models that, given rules, will generate new data) can be helpful to initiate the investigation of questions that have not received (or perhaps cannot receive) extensive empirical treatment, or for which usual statistical approaches fall short. In particular, we argue that computational approaches can serve a dual purpose. First, they can deliver a more predictive science, because they are explicitly data-driven. Second, they can guide the attention of researchers onto mechanisms of interest; in a context where time and resources are finite, and the urgency to understand ecological systems is high, this may be the main selling point of computational techniques.

In a thought-provoking essay, Markowetz (2017) suggests that *all biology is computational biology* - the rationale behind this bold statement being that integrating computational advances, novel mathematical tools, and the usual data from one field, has a high potential to deliver synthesis. A more reasonable statement would be that *all ecology can benefit from computational ecology*, as long as we can understand how it interacts with other approaches; in this paper, we attempt to situate the practice of computational ecology within the broader scope of ecological research. The recent years have given us an explosion of new tools, training opportunities, and mechanisms for data access. One can assume that computational approaches will become more tempting, and more broadly adopted. This requires us to address the questions of the usefulness and promises of this line of research, as well as the caveats associated with it. In particular, we highlight the ways in which computational ecology differs from, and complements, ecological modelling that does not involve data directly. We finally move on to the currency of collaborations between different sub-disciplines of ecologists, and discuss the need to add more quantitative skills in ecological training.

Advancing ecology through computational techniques is an ongoing work, and already delivered many results (some of which we discuss in the text). To elevate this approach, the community of practising ecologists needs to establish baselines of appropriate practices for the sharing and re-use of existing data, especially when they are massively aggregated and re-purposed; reach a consensus on a common core of training which enables students to explore computational approaches in addition to more usual approaches. Ultimately, a better integration of computational techniques in the practice of ecological research has the potential to improve transparency and reproducibility, and facilitate the synthesis of ecological knowledge.

## A success story: Species Distribution Models

The practice known as “species distribution modelling” (and the species distribution models, henceforth SDMs, it generates) is a good example of computational practices generating novel ecological insights. At their core, SDMs seek to model the presences or absences of a species based on previous observations of its presences or absences, and knowledge of the environment in which the observations were made. More formally, SDMs can be interpreted as having the form P(S|E) (or P(S = 1|E) for presence-only models), where S denotes the presence of a species, and E is an array of variables representing the local state of the environment at the point where the prediction is made (the location is represented, not by its spatial positions, but by a suite of environmental variables).

As Franklin (2010) highlights, SDMs emerged at a time where access to computers *and* the ability to effectively program them became easier. Although ecological insights, statistical methods, and data already existed, the ability to turn these ingredients into something predictive required what is now called “computational literacy” - the ability to abstract, and automate, a system in order to generate predictions through computer simulations and their validation. One of the strengths of SDMs is that they can be used either for predictions or explanations of where a given species occur (Elith & Leathwick 2009) and can be corroborated with empirical data. To calculate P(S|E) is to make a prediction (what are the chances of observing species S at a given location), that can be refined, validated, or rejected based on cross-validation (Hijmans 2012) or de *novo* field samplig (West et al. 2016). To understand E, *i.e.* the environmental aspects that determine species presence, is to form an explanation of a distribution that relates to the natural history of a species.

SDMs originated as statistical and correlative models, and are now incorporating more ecological theory (Austin 2002) - being able to integrate (abstract) ideas and knowledge with (formal) statistical and numerical tools is a key feature of computational thinking. In fact, one of the most recent and most stimulating developments in the field of SDMs is to refine their predictions not through the addition of more data, but through the addition of more processes (Franklin 2010). These SDMs rely on the usual statistical models, but also on dynamical models (for example simulations; e.g. Wisz et al. (2012) or Pellissier et al. (2013) for biotic interactions, and Miller & Holloway (2015) for movement and dispersal). What they lack in mathematical expressiveness (i.e. having a closed-form solution (Borwein & Crandall 2013), which is often ruled out by the use of stochastic models or agent-based simulations), they assume to gain in predictive ability through the explicit consideration of more realistic ecological mechanisms (D’Amen et al. 2017; Staniczenko et al. 2017).

SDMs have been a success, but there are many other areas of ecology that could be improved by a marriage of computational ecology and empirical data. The novel use of genomic RNA-seq data and existing worldclim climate data allowed creating random forest models in order to make predictions where yellow warbler populations, a species of conservation concern, are most vulnerable to climate change (Bay et al. 2018). Environmental DNA metabarcoding data coupled with machine learning approaches and linear models was used to create, test, and predict biodiversity indices for benthic foraminifera, which can be applied to monitoring health of fish farm ecosystems (Cordier et al. 2017). The increase in data volume, coupled with access to computing techniques and power, will result in a multiplication of these boundary-pushing studies in the next years.

## 2. Outlining computational ecology

Most research approaches exist on a gradient. In this section, we will outline research practices which differ enough in their approaches to fall under the umbrella of computational science, and specifically discuss how they can provide novel information. We will first show how computational ecology complements other research approaches, then discuss how it can be used in the current context to facilitate interactions between theoretical and empirical research.

### 2.1. Computational ecology in focus

The specific example of predator-prey interactions should be a familiar illustration of how the same problem can be addressed through a variety of research approaches (fig. 1). The classic predator-prey equations of Lotka & Volterra are an instance of a “modelling” based perspective, wherein mathematical analysis reveals how selected parameters (rates of interactions and growth) affect an ecologically relevant quantity (population stability and coexistence). These models, although they have been formulated to explain data generated through empirical observations, are disconnected from the data themselves. In fact, this family of model lies at the basis of a branch of ecological modelling that now exists entirely outside of data (Ackland & Gallagher 2004; Gyllenberg et al. 2006; Coville & Frederic 2013). These purely mathematical models are often used to describe trends in time series. But not all of them hold up to scrutiny when explicitly compared to empirical data. Gilpin (1973) famously reports that based on the predictions of the Lotka-Volterra model, hares in the Hudson bay are feeding on Lynx - this example goes to show that blindly applying models is dangerous, and their output should be framed in the context of external data.

**Figure 1.**
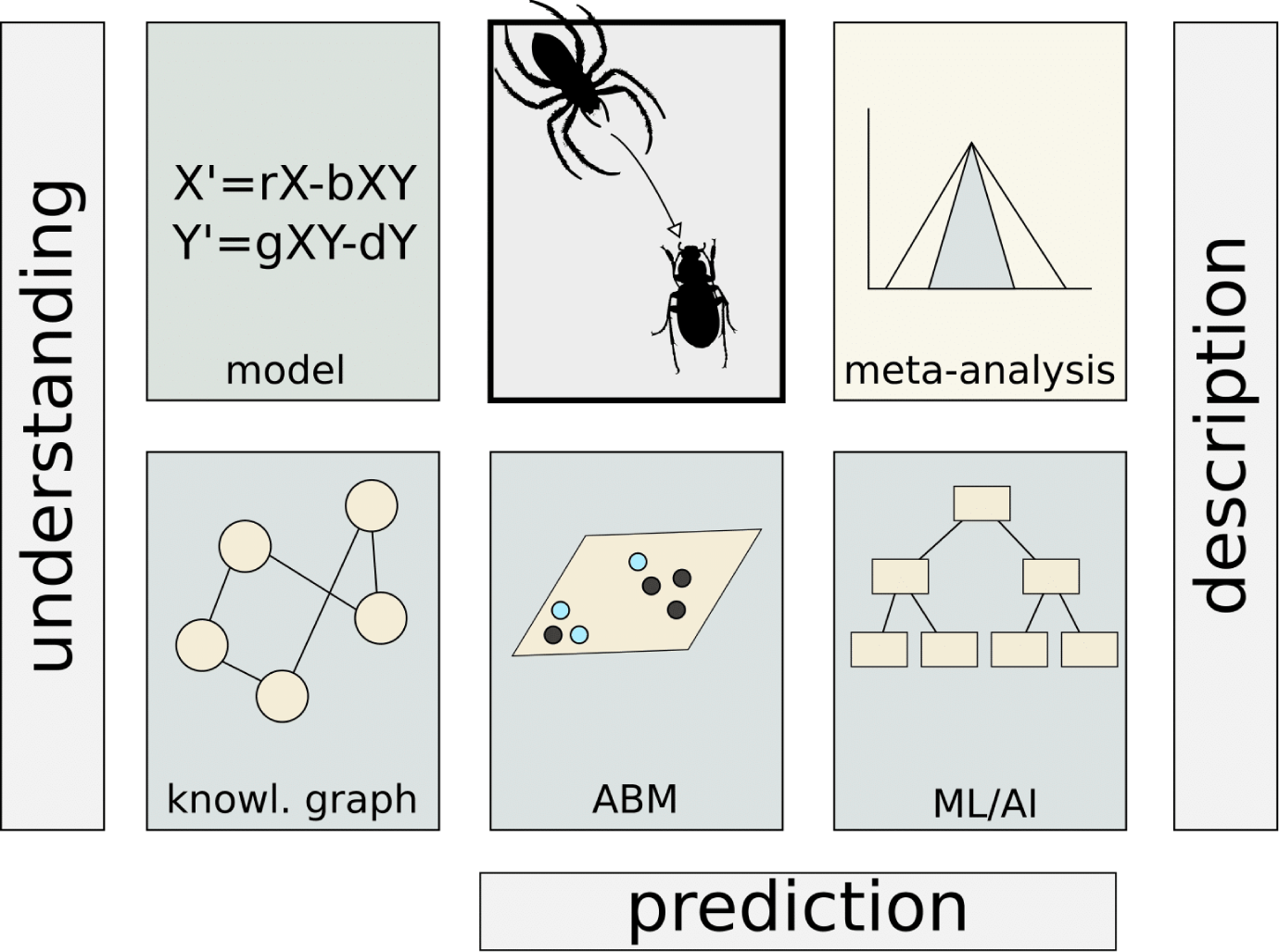
An overview of how computational approaches can complement other research approaches. On the top line, we have represented empirical studies (center) as well as modelling (left) and meta-analysis (right; represented as a funnel plot) approaches. In the bottom line, we have represented three possible approaches to study predator-prey relationships: knowledge graphs can represent interactions between the concepts; agent-based modelling can provide some predictions about the future of the system; methods from machine learning can assist both in understanding and prediction. Importantly, the goal of these approaches should always be to return to empirical data.

By contrast Sallan et al. (2011) study the same issue (sustained persistence and fluctuations of predator-prey couples through time) using a paleo-ecological timeseries, and interpret their data in the context of predictions from the Lotka-Volterra family of models (namely, they find support for Lotka-Volterra-like oscillations in time). Although dynamical models and empirical data interact in this example, they do not do so directly; that is, the analysis of empirical data is done within the context of a broad family of model, but not coupled to *e.g.* additional simulations. The two are done *in parallel*, and not so much *in interaction.* A number of other models have been shown to generate predictions that quantitatively match empirical data (Nicholson & Bailey 1935; Beverton & Holt 1957) - this represents, in our opinion, the sole test of whether a mathematical model is adapted to a particular problem and system. While models are undeniably useful to make mechanisms interact in a low-complexity setting, it is a grave mistake to assume they will, in and of themselves, be relevant to empirical systems.

Meta-analyses, such as the one by Bolnick & Preisser (2005), are instead interested in collecting the outcome of observational and manipulative studies, and synthesizing the *effects* they report. These are often purely *statistical*, in that they aggregate significance and effect size, to measure how robust a result is across different systems. Meta-analyses most often require a *critical mass* of pre-existing papers (Lortie et al. 2013). Although they are irreplaceable as a tool to measure the strength of results, they are limited by their need for primary literature with experimental designs that are similar enough.

Predator-prey (and other biotic) interactions have been studied with a few computational approaches to date. Colon et al. (2015) show how an agent-based model can guide the interpretation of the same system represented as ordinary differential equations. This is an important result, as it offers suggestions to bridge families of models - not only can agent-based approaches provide answers about the biological systems of interest, they can also provide information about the behaviour of other families of models. Although this example is primarily model-driven, there are a number of data-driven approaches that rely on computational techniques. One example is the prediction of species interactions. Stock et al. (2017) suggested linear filtering to identify false-negatives (*i.e.* interactions that exist, but may have been missed) in empirical dataset. This can guide sampling in the field, and is to an extent a predictive task, but cannot inform our understanding of the system. Similarly, Desjardins-Proulx et al. (2017a) used various recommender systems to infer the prey items of predators based on knowledge of (i) diet and (ii) functional traits. This results in testable predictions, but is not necessarily increasing our understanding of the rules involved in the system.

Chen et al. (2016) used symbolic regression to infer a differential equations model *from data* about predator-prey interactions. This is a fascinating result, as it shows just how much signal is contained in data: enough to describe a mathematical model explaining their behaviour. And while understanding mechanisms by looking at a time series may be difficult, understanding the mechanisms when studying equations dictated by the data themselves is feasible. In a similar vein, Desjardins-Proulx et al. (2017b) suggest that logic networks, which describe the relationships between concepts, can be inferred by optimizing a knowledge bank on the data. This category of approaches offer the opportunity to increase our understanding of empirical data, not by thinking deeply about the rules, but by extracting the rules from the data.

### 2.2. Computational ecology in context

In *Life on the Mississippi*, Mark Twain wrote that “There is something fascinating about science. One gets such wholesale returns of conjecture out of such a trifling investment of fact”. This is a good description of the purpose of computational ecology: in a data-limited context, merging phenomenological models with pre-existing datasets is a way to efficiently develop conjectures, or more appropriately, build on our knowledge of models and data to put forward testable, quantified hypotheses. Perretti et al. (2013) intriguingly report that model-free inference based on data *always* outperforms the best model: in other words, we do not understand ecological systems as well as we think, and approaches putting the data first might always outperform those relying on expert knowledge. Pascual (2005) outlined that computational ecology has a unique ability to go from the complex (natural systems) to the simple (representations and conceptual models), and back (testable predictions). Although the natural world is immensely complex, it is paradoxically the high degree of model abstraction in computational approaches that gives them generality across several systems. In the years since this article was published, the explosion in machine learning tools and their predictive ability, and their adoption by ecologists (Thessen 2016) should have changed the situation quite significantly.

Yet, with the exception of a still narrow family of problems that can be addressed by remote-sensing or meta-genomics, there has been no regime shift in the rate at which ecological data are collected. Observations from citizen science accumulate, but are highly biased by societal preferences rather than conservation priority (Donaldson et al. 2016; Troudet et al. 2017), by proximity to urban centers and infrastructure (Geldmann et al. 2016), *as well as* by the interaction between these factors (Tiago et al. 2017). In addition, Lindenmayer & Likens (2018) raise the significant concern that the “culture” of ecology must be maintained - even in the context of a sudden (though debatable) avalanche of data, ecology as a field should always put robust hypotheses *first.* This is especially true since our needs for testable and actionable predictions increased dramatically. This provides a clear mission statement for computational ecology: refining the models and further integrating them with data is necessary, and using methods that work well on reduced amounts of heterogeneous data must be part of this effort. Enthusiastic reports about the big data revolution coming to ecology (Hampton et al. 2013; Soranno & Schimel 2014) have been premature at best, and the challenge associated with most of our datasets being decidedly *tiny* cannot be easily dismissed.

Yet data, even small, are “unreasonably effective” (Halevy et al. 2009) - they can reveal trends and signal that may not be immediately apparent from causal modelling alone, for example. Ecological models make, by definition, high accuracy predictions, but they tend to be difficult to test (Rykiel 1996) - models relying on precise mathematical expressions can be difficult to calibrate or parameterize. Observations (field sampling) or manipulative approaches (micro/meso/macro-cosms, field experiments) are highly accurate (but have also immense human and monetary costs that limit the scale at which they can be applied). There is simply too much nature around for us to observe, monitor, and manipulate it all. In this perspective, computational approaches able to generalize some rules from the data (Desjardins-Proulx et al. 2017a, 2017b) may help guide the attention of researchers onto mechanisms that are worthy of a deeper investigation. Computational approaches will more likely shine *in support* to more established areas of research.

Recent advances in computational epidemiology (reviewed in Marathe & Ramakrishnan 2013) provide an interesting roadmap for computational ecology: there have been parallel advances in (i) adapting data acquisition to maximize the usefulness of novel data analyses methods, (ii) integration of novel analytical methods from applied mathematics *and* social sciences, mostly related to computations on large graphs, to work on pre-existing data, and (iii) a tighter integration of models to data fluxes to allow near real-time monitoring and prediction. All of these things are possible in ecological research. In fact, recent examples (Bush et al. 2017; Harris et al. 2017; Dietze et al. 2018; White et al. 2018) suggest that near real-time forecasting of biodiversity is becoming feasible, and is identified by computational ecologists as a key priority.

## 3 En route towards synthesis

Ecological synthesis, usually defined as the integration of data and knowledge to increase scope, relevance, or usability of results both across and within sub-fields (Carpenter et al. 2009), is an essential first step in order to achieve policy relevance (Baron et al. 2017). Most of the global policy challenges have an ecological or environmental component, and outside of the socio-ecological, socio-economical, socio-cultural, aspects, ecologists can contribute to the mitigation or resolution of these challenges by i) assessing our knowledge of natural systems, ii) developing methods to produce scenarios using state-of-the-art models and tools, and iii) communicating the output of these scenarios to impact policy-making. White et al. (2015) propose that this falls under the umbrella of *action ecology, i.e.* using fundamental knowledge and ecological theory to address pressing, real-world questions.

Raghavan et al. (2016) suggest that this approach can also accommodate stakeholder knowledge and engagement. By building models that rely on ecological concepts, empirical data, and stakeholder feedback, they propose a *computational agroecology* program, to use computational tools in the optimization of sustainable agricultural practices. This example suggests that not only can computational approaches yield fundamental research results in a short time frame, they can also be leveraged as a tool for applied research and knowledge transfer now. The definition of “a short time” is highly sensitive to the context - some predictions can be generated using routine tools (in a matter of weeks), whereas some require to develop novel methodologies, and may require years. Accelerating the time to prediction will, in large part, require the development of software that can be deployed and run more rapidly. Computational ecology is nevertheless nimble enough that it can be used to iterate rapidly over a range of scenarios, to inform interactions with policy makers or stakeholders in near real time. We need to mention that there is a lower bound on time to prediction: some applications require different degrees of accuracy. While an approximate result is good enough for fundamental research, outputs used to enact policy making that can affect thousands of citizens (and change the dynamics of a region or an ecosystem) require a better accuracy. The variety of computational techniques allows moving across these scales, while the advances in programming practices and computing power decreases the severity of the accuracy/runtime tradeoff.

### 3.1. Mapping the domains of collaboration

Understanding how computational ecology will fit within the broader research practices requires answering three questions: what can computational ecology bring to the table, what are the needs of computational ecologists, and what are the current limitations of computational approaches that could limit their immediate applicability. It seems, at this point, important to minimize neither the importance nor the efficiency of sampling and collection of additional data. Sampling is important because ecological questions, no matter how fundamental, ought to be grounded in phenomena happening in nature, and these are revealed by observation or manipulation of natural systems. Sampling is efficient because it is the final arbiter: how good any prediction is at explaining aspects of a particular empirical system is determined by observations of this system, compared to the predictions.

Relying heavily on external information implies that computational research is dependent on standards for data representation. The Ecological Metadata Language (Fegraus et al. 2005) is an attempt at standardizing the way meta-data are represented for ecological data; adherence to this standard, although it has been shown to improve the ease of assembling large datasets from single studies (Gil et al. 2011), is done on a voluntary basis (and its uptake is therefore abysmal). An alternative approach is to rely on community efforts to pre-curate and pre-catalog ecological data, such as with the flagship effort *EcoDataRetriever* (Morris & White 2013). Yet even this approach is ultimately limited because of the human factor involved — when the upstream data change, they have to be re-worked into the software. A community consensus on data representation, although unlikely, would actually solve several problems at once. First, it would make the integration of multiple data sources trivial. Second, it would provide clear guidelines about the input and storage of data, thus maybe improving their currently limited longevity (Vines et al. 2014).

Finally, it would facilitate the integration of data and models with minimum efforts and risk of mis-communication, since the format would be the same for all. To this extent, a recent proposal by Ovaskainen et al. (2017) is particularly interesting: rather than deciding on formats based on knowledge of eco-informatics or data management best practices, why not start from the ecological concepts, and translate them in digital representation? The current way to represent e.g. biodiversity data has largely been designed based on the needs of collection managers, and bears little to no relevance to most extant research needs. Re-designing the way we store and manipulate data based on research practices is an important step forward, and will ultimately benefit researchers. To be generalized, this task requires a strong collaboration between ecologists with topic expertise, ecologists with field expertise, and those of us leaning closest to the computational part of the field.

With or without a common data format, the problem remains that we have very limited insights into how error in predictions made on synthetic datasets will propagate from an analysis to another (Poisot et al. 2016); in a succession of predictive steps, do errors at each step amplify, or cancel one another? Biases exist in the underlying data and in the models used to generate the predictions, and these biases can manifest in three possible outcomes. First, predictions from these datasets accumulate bias and cannot be used. Second, because the scale at which these predictions are expressed is large, errors are (quantitatively) small enough to be over-ridden by the magnitude of actual variation. Finally, in the best-case but low-realism scenario, errors end up cancelling each other out. The best possible way to understand how errors propagate is to validate predictions *de novo*, through sampling. Model-validation methods can be used, as they are with SDMs (Hijmans 2012), but *de novo* sampling carries the additional weight of being an independent attempt at testing the prediction. Improved collaborations on this aspect will provide estimates of the robustness of the predictions, in addition to highlighting the steps of the process in which uncertainty is high — these steps are natural candidates for additional methodological development.

Finally, there is a need to assess how the predictions made by purely computational approaches will be fed back into other types of research. This is notably true when presenting these approaches to stakeholders. One possible way to make this knowledge transfer process easier is to be transparent about the way predictions were derived: which data were used (with citations for credits and unique identifiers for reproductibility), which software was used (with versions numbers and code), and what the model / simulations do (White et al. 2013). In short, the onus is on practitioners of computational research to make sure we provide all the information needed to communicate how predictions came to be.

### 3.2. Establishing the currencies of collaboration

An important question to further the integration of computational approaches to the workflow of ecological research is to establish *currencies* for collaborations. Both at the scale of individual researchers, research group, and larger research communities, it is important to understand what each can contribute to the research effort. As ecological research is expected to be increasingly predictive and policy-relevant, and as fundamental research tends to tackle increasingly refined and complex questions, it is expected that research problems will become more difficult to resolve. This is an incentive for collaborations that build on the skills that are specific to different approaches.

In an editorial to the *New England Journal of Medicine*, Longo & Drazen (2016) characterized scientists using previously published data as “research parasites” (backlash by a large part of the scientific community caused one of the authors to later retract the statement - Drazen (2016)). Although community ecologists would have, anyways, realized that the presence of parasites indicates a healthy ecosystem (Marcogliese 2005; Hudson et al. 2006), this feeling of unfair benefit for ecological data re-analysis (Mills et al. 2015) has to be addressed, because it has no empirical support. The rate of data re-use in ecology is low and has a large delay (Evans 2016), and there are no instances of re-analysing existing data for the same (or similar) purpose they were produced for. There is a necessary delay between the moment data are available, and the moment where they are aggregated and re-purposed (especially considering that data are, at the earliest, published at the same time as the paper). This delay is introduced by the need to understand the data, see how they can be combined, develop a research hypothesis, etc..h

On the other hand, there are multiple instances of combining multiple datasets collected at different scales to address an entirely different question (see GBIF 2016 for an excellent showcase) - it is more likely that data re-use is done with the intent of exploring different questions. It is also worth remembering that ecology as a whole, and macroecology and biogeography in particular, already benefit immensely from data re-use. For example, data collected by citizen scientists are used to generate estimates of biodiversity distribution, but also set and refine conservation targets (Devictor et al. 2010); an overwhelming majority of our knowledge of bird richness and distribution comes from the *eBird* project (Sullivan et al. 2009, 2014), which is essentially fed by the unpaid work of citizen scientists.

With this in mind, there is no tip-toeing around the fact that computational ecologists will be *data consumers*, and this data will have to come from ecologists with active field programs (in addition to government, industry, and citizens). Recognizing that computational ecology *needs* these data as a condition for its continued existence and relevance should motivate the establishment of a way to credit and recognize the role of *data producers* (which is discussed in Poisot et al. 2016, in particular in the context of massive dataset aggregation). Data re-users must be extremely pro-active in the establishment of crediting mechanisms for data producers; as the availability of these data is crucial to computational approaches, and as we do not share any of the cost of collecting these data, it behooves us to make sure that our research practices do not accrue a cost for our colleagues with field or lab programs. Encouraging conversations between data producers and data consumers about what data will be shared, when, and how databases will be maintained will improve both collaborations and research quality. In parallel, data producers can benefit from the new analytical avenues opened by advances in computational ecology. Research funders should develop financial incentives to these collaborations, specifically by dedicating a part of the money to developing and implementing sound data archival and re-use strategies, and by encouraging researchers to re-use existing data when they exist.

### 3.3. Training data-minded ecologists

The fact that data re-use is not instantaneously convenient reveals another piece of information about computational ecology: it relies on different skills, and different tools than those typically used by field ecologists. One of the most fruitful avenues for collaboration lies in recognizing the strengths of different domains: the skills required to assemble a dataset (taxonomic expertise, natural history knowledge, field know-how) and the skills required to develop robust computational studies (programming, applied mathematics) are different. Because these skills are so transversal to any form of ecological research, we are confident that they can be incorporated in any curriculum. If anything, this calls for increased collaboration, where these approaches are put to work in complementarity.

Barraquand et al. (2014) highlighted the fact that professional ecologists received *less* quantitative and computational thinking that they think should be necessary. Increasing the amount of such training does not necessarily imply that natural history or field practice will be sacrificed on the altar of mathematics: rather, ecology would benefit from introducing more quantitative skills and reasoning across all courses, and introductory ones in particular (Hoffman et al. 2016). Instead of dividing the field further between empirically and theoretically minded scientists, this would showcase quantitative skills as being transversal to all questions that ecology can address. What to teach, and how to integrate it to the existing curriculum, does of course require discussion and consensus building by the community.

A related problem is that most practising ecologists are terrible role models when it comes to showcasing good practices of data management (because there are no incentives to do this); and data management is a crucial step towards easier computational approaches. Even in the minority of cases where ecologists do share their data on public platforms, there are so few metadata that not being able to reproduce the original study is the rule (Roche et al. 2014, 2015). This is a worrying trend, because data management affects how easily research is done, regardless of whether the data are ultimately archived. Because the volume and variety of data we can collect tends to increase over time, and because we expect higher standards of analysis (therefore requiring more programmatic approaches relying on the use or development of purpose-specific code), data management has already became a core skill for ecologists to acquire.

This view is echoed in recent proposals. Mislan et al. (2016) suggested that highlighting the importance of code in most ecological studies would be a way to bring the community to adopt higher standards, all the while de-mystifying the process of producing code. As with increased mandatory data release alongside more reproducible publication required by funding agencies, mandatory code release would benefit a more reproducible science and show how data were transformed during the analysis. This also requires teaching ecologists how to evaluate the quality of the software they use (Poisot 2015). Finally, Hampton et al. (2015) proposed that the “Tao of Open Science” would be particularly beneficial to the entire field of ecology; as part of the important changes in attitude, they identified the solicitation and integration of productive feedback throughout the research process. Regardless of the technical solution, this emphasizes the need to foster, in ecologists in training, a culture of discussion across disciplinary boundaries.

All of these points can be distilled into practical training recommendations for different groups in the community of ecologists. Classes based around lab or field experience should emphasize practical data management skills which have been validated as best practices by the community (Soyka et al. 2017), and introduce tools that would make the maintenance of data easier. Modelling classes, especially when concerned about purely mathematical models, should add modules on the way these models can be integrated with empirical data. Finally, computational classes should emphasize communication skills: what do these new tools do, and how can they be used by other fields in ecology; but also, how do we properly track citations to data, and give credit to data producers? Building these practices into training would ensure that the next generation of ecologists will be able to engage in a meaningful dialogue across methodological boundaries.

## 4 Concluding remarks

None of the theoretical, mathematical, computational approaches to ecological research have any intrinsic superiority - in the end, direct observation and experimentation trumps all, and serve as the validation, rejection, or refinement of predictions derived in other ways, but lacks the scaling power to be the only viable solution. The growing computational power, growing amount of data, and increasing computational literacy in ecology means that producing theory and predictions is becoming cheaper and faster (regardless of the quality of these products). Yet the time needed to test any prediction is not decreasing (or at least not as fast). Computational science has resulted in the development of many tools and approaches that can be useful to ecology, since they allow ecologists of all kinds to wade through these predictions and data. Confronting theoretical predictions to data is a requirement, if not the core, of ecological synthesis; this is only possible under the conditions that ecologists engage in meaningful dialogue across disciplines, and recognize the currencies of their collaborations.

Discussing the place of computational ecology within the broader context of the ecological sciences will highlight areas of collaborations with other areas of science. Thessen (2016) makes the point that long-standing ecological problems would benefit from being examined through a variety of machine learning techniques - We fully concur, because these techniques usually make the most of existing data (Halevy et al. 2009). Reaching a point where these methods are routinely used by ecologists will require a shift in our culture: quantitative training is currently perceived as inadequate (Barraquand et al. 2014), and most graduate programs do not train ecology students in contemporary statistics (Touchon & McCoy 2016).

Ultimately, any additional data collection has its scope limited by financial, human, and temporal constraints — or in other words, we need to chose what to sample, because we can’t afford to sample it all. Computational approaches, because they can work through large amounts of data, and integrate them with models that can generate predictions, might allow answering an all important question: what do we sample, and where? Some rely on their ecological intuition to answer; although computational ecologists may be deprived of such intuitions, they have the know-how to couple data and models, and can meaningfully contribute to this answer. Computational ecology is also remarkably cost-effective. Although the reliance on advanced research computing incurs immense costs (including hardware maintenance, electrical power, and training of highly qualified personnel; these are often absorbed by local or national consortia), it allows the generation of predictions that are highly testable. Although the accuracy of these predictions is currently unknown (and will vary on a model/study/question basis), any additional empirical effort to *validate* predictions will improve their quality, reinforcing the need for dialogue and collaborations.

## Acknowledgements

TP thanks Dr. Allison Barner and Dr. Andrew McDonald for stimulating discussions, and the Station de Biologie des Laurentides de l’Universite de Montreal for hosting him during part of the writing process. TP thanks the Canadian Institute for Ecology and Evolution for financial support. We thank the volunteers of Software Carpentry and Data Carpentry, whose work contribute to improving the skills of ecologists. Carabid picture by Maxime Dahirel (CC-BY 4.0), spider image by Sidney Frederic Harmer, Arthur Everett Shipley, digitized by Maxime Dahirel (CC-BY 4.0).

